# Unraveling AMPK and BET regulation of immune checkpoint biology: implications for personalized medicine

**DOI:** 10.64898/2026.01.26.701869

**Authors:** Christina S. Ennis, Kunlin Huang, Allison N. Casey, Michael Seen, Gerald V. Denis

## Abstract

Triple negative breast cancer (TNBC) patients with comorbid Type 2 diabetes (T2D) show worse survival compared to nondiabetic TNBC patients. Immune checkpoint blockade (ICB) has unclear benefit in TNBC. Immune suppression in T2D, and use of metformin, an activator of 5’ Adenosine Monophosphate-activated Protein Kinase (AMPK), in such patients, prompted us to examine AMPK regulation of immune checkpoint expression. Improved ICB efficacy may optimize outcomes for certain TNBC patients. We have also been exploring the role of Bromodomain and ExtraTerminal domain (BET) proteins (BRD2, BRD3, BRD4) in regulation of checkpoint molecules in immune cell subsets, including CD4+, CD8+ T cells, and NK cells. BET proteins are important transcriptional co-regulators, critical for proliferation and metastasis in many cancer types, including TNBC. We observed differential BET regulation of immune checkpoint proteins, specifically TIM-3, TIGIT, PD-1 and CTLA-4, on αCD3/αCD28-stimulated peripheral blood mononuclear cells by flow cytometry. Chemical inhibition of AMPK with Compound C, and with the pan-BET inhibitor JQ1 or the BRD4-selective PROTAC inhibitor MZ-1, revealed that BET proteins regulate PD-1 and CTLA-4 through an AMPK-dependent pathway and TIM-3 and TIGIT through an AMPK-independent pathway. Personalized approaches to ICB treatment of TNBC patients with comorbid T2D should improve outcomes.

## Introduction

Immune evasion is one of several strategies that solid tumors are known to take as they interact with the tumor microenvironment and evolve into more aggressive and pro-metastatic states.^1^ These mechanisms enable malignant clones to circumvent T cell mediated anti-tumor immunity through the induction of T cell exhaustion.^2^ This altered differentiation state, commonly observed during periods of persistent antigen exposure, is characterized by the metabolic impairment and functional weakening of T and NK cell anti-tumor activity.^3^ Consequently, patients with exhausted immune cells become more susceptible to the establishment and progression of cancer. Progress has been made towards reversing immune exhaustion via immune checkpoint blockade (ICB), which block critical ligand-receptor interactions,^4^ such as PD-1/PD-L1, CTLA-4/CD80/CD86, TIGIT/CD155 and TIM-3/GAL9. This signaling triggers internal signaling cascades that establish the exhausted state. However, despite its promise, a significant proportion of patients fail to respond to ICB therapy.^5^ Understanding the mechanisms underlying this response remains a pressing challenge in the field. A particularly promising avenue for investigation lies in the exploration of the transcriptional networks regulating the expression of immune checkpoint (IC) genes.^6^ Unraveling these networks may yield insights that could be harnessed to improve therapeutic outcomes for cancer patients who being treated with ICB approaches.

We have been investigating triple negative breast cancer (TNBC), the most aggressive type of breast cancer, which has required new targeted therapies because of its low or absent expression of estrogen receptor (ER), progesterone receptor and epidermal growth factor receptor 2. TNBC exhibits significant tumoral heterogeneity that associates with treatment resistance and recurrence, including in the tumor immune microenvironment.^7^ TNBC has been considered as potentially amenable to ICB approaches, including anti-PD-1 (pembrolizumab and nivolumab), anti-PD-L1 (atezolizumab, avelumab and durvalumab), and anti-CTLA-4 (tremelimumab) monoclonal antibody combinations, for patients with adequate expression of these surface markers. Although these agents are typically well tolerated, efficacy in TNBC is still less than optimal, and some clinical trials have been terminated without encouraging outcomes.^8^ Improved understanding of IC gene regulation could help improve TNBC patient responses to ICB.

The bromodomain and extraterminal (BET) family of transcriptional co-regulators, including BRD2, BRD3, and BRD4, are well established as key players in cancer biology, inflammation^9^ and anti-tumor immunity.^10^ BET proteins are epigenetic transcriptional co-regulators recognize and bind acetyl groups on nucleosomal histones, interactions that are amenable to small molecule inhibition.^11^ BET proteins contain highly conserved bromodomain motifs that interact with acetylated histones, facilitating the recruitment of transcriptional regulators and controlling the expression of multiple critical genes.

Aberrant regulation of BET proteins has been implicated in cancer metastasis and progression, making them attractive targets for cancer therapy. Inhibition of BRD4, for example, has shown promising effects in suppressing TNBC,^12,13^ including in combination with CDK4/6 inhibitors.^14^ However, we have also established in TNBC that individual BET proteins regulate independent transcriptional networks that differ from each other and may oppose each other, depending on the context.^15^ Therefore, strategies that rely on pan-BET inhibitors in TNBC, such as JQ1, ignore these differences and may promote undesirable and dangerous biological outcomes.^16^

Furthermore, we and others have established that BET proteins are essential regulators of PD-1 ^17,18^ and PD-L1 ^17,19^ transcription. Experiments with siRNA knockdown reveal that individual BET proteins, specifically BRD2 and BRD4, but not BRD3, independently regulate PD-1 gene expression. Higher levels of BRD2 or BRD4 in individual patient samples associate with higher expression of PD-1, suggesting avenues for personalized approaches to improve anti-tumor immunity in TNBC.^17^ There is growing evidence that combining BET inhibition with ICB has efficacy for certain solid tumors,^19-21^ as well as ICB resistant melanoma.^22^

In addition to the BET transcriptional network, 5’ Adenosine Monophosphate-activated Protein Kinase (AMPK) has also been implicated in cancer progression and immune function in cancer,^23^ including regulation of PD-1 expression.^24^ AMPK serves as a master regulator of cellular energy balance, responding to energy stress by restoring equilibrium through the modulation of anabolic and catabolic processes. AMPK is a heterotrimeric protein complex involved in diverse metabolic pathways, including lipid metabolism, glucose transport, and uptake.^25^ AMPK activation has been linked to the development of cancer, promoting the formation of mammospheres in breast cancer.^26^ Recent studies have demonstrated the importance of AMPK in regulating the epigenome of cancer cells.^27^ Knockdown of BRD2 specifically has been shown to increase fatty acid oxidation in pancreatic β cells, implicating BET proteins in the regulation of AMPK-dependent processes^28^ and suggesting that joint targeting of AMPK and BET proteins may have efficacy for certain cancers.^29^ These findings underscore the interconnectedness of these central pathways in cancer biology, highlighting their potential as key players in the development of novel therapeutic strategies.

In the present study, we aimed to investigate how BET proteins regulate IC genes and whether the AMPK signaling pathway is involved in this regulation. Using peripheral blood mononuclear cells (PBMCs) from healthy adult donors treated with various inhibitors of either BET or AMPK, we demonstrated that BET family members have a differential regulation on the expression of IC genes, with separate signaling pathways likely converging on BRD2/BRD3/BRD4 to act independently as effectors at each promoter. Notably, we observed an AMPK-dependent regulation of PD-1 and CTLA-4 that is differentially regulated from TIGIT and TIM-3. These findings shed light on the intricate molecular mechanisms underlying ICs and suggest innovative metabolic-based approaches to prevent immune exhaustion or overcome ICB resistance.

## Materials and Methods

### PBMC Isolation

Patient leukopaks were obtained commercially from New York Biologics Blood Banking, isolated by leukapheresis from different, normal adult donors. Peripheral blood mononuclear cells (PBMCs) were isolated by Ficoll-Paque™ PLUS (GE Healthcare) centrifugation. Cell yields and viability were determined via hemocytometer. Freshly isolated PBMCs were resuspended and cryopreserved in 90% fetal bovine serum (FBS, Sigma-Aldrich) and 10% dimethyl sulfoxide (DMSO, ATCC) at -80°C, then transferred to liquid nitrogen for long-term cryopreservation. The cryopreserved cells could be thawed at any point weeks or months later, removed from the DMSO freezing medium, diluted into fresh RPMI 1640 Medium (1x) (Gibco, ThermoFisher) with 10% FBS and stimulated or expanded, with >90% viability. To obtain T cell subsets for proliferation and cytokine production assays, CD8+ T cells were purified from leukopaks by negative selection with a specialized isolation kit (Miltenyi 130-096-495) and magnetic depletion of non-CD8+ T cells as we have previously reported.

### Ex vivo T cell activation

Plates were prepared with 10 μg/ml anti-human CD3 antibody (Clone OKT3, Ultra-LEAF, BioLegend) and incubated at 37°C for an hour to ensure antibody binding to the plate surface. PBMCs were thawed and resuspended with pre-warmed RPMI1640 supplemented with 2 mM L-glutamine, penicillin+streptomycin, 50 µM 2-mercaptoethanol, 10% FBS, and containing 2 μg/ml anti-human CD28 antibody (Clone CD28.2, Ultra-LEAF, BioLegend), then layered onto the treated plates. Pan-BET inhibitor JQ1 (Tocris), BRD4-selective PROTAC degrader MZ-1 (Tocris), and AMPK inhibitor Compound C (Tocris) were dissolved in DMSO to final concentrations of 400 nM, 50 nM and 5 µM respectively. These stock solutions were kept at -20°C until use, diluted to the final concentration in media, then added to cells immediately. Cells were incubated for 48-72 hours at 37°C in a 5% CO_2_ environment with 100% humidity.

### Flow Cytometry Analysis

Cell surface staining was performed with monoclonal antibodies: CD3 Alexa Fluor 488 (BioLegend, Clone OKT3), CD4 APC (BioLegend, Clone SK3), CD8 PerCp/Cy5.5 (BioLegend, Clone SK1), CD56 BUV737 (BD Biosciences, Clone NCAM16.2 (RUO)), TIM-3 BV421 (BioLegend, Clone F38-2E2), TIGIT BV605 (BioLegend, Clone A15153G), PD-1 PE (BioLegend, Clone EH12.2H7), and CTLA-4 PE/Cy7 (BioLegend, Clone L3D10). Zombie NIR™ Fixable Viability Dye (BioLegend) was used to identify live cells. Brilliant Stain Buffer (BD Biosciences) was used to reduce nonspecific fluorescent dye interactions, and True-Stain Monocyte Blocker (BioLegend) was used to reduce non-specific binding of monocytes. Signals from at least 100,000 PBMCs per sample were measured on an LSR II (BD Biosciences) flow cytometer in the BUMC Flow Cytometry Core Facility. Data analysis was performed with FlowJo software (FlowJo LLC, USA).

For intracellular cytokine analysis, PBMCs or purified, untouched CD8+ T cells were stimulated for 48 hrs with Cell Stimulation Cocktail™ (eBioscience, Invitrogen). Brefeldin A (eBioscience, Invitrogen) was added for the last 24 hrs to allow for intracellular cytokine accumulation. Cells were then fixed, permeabilized, and stained with monoclonal antibodies: CD4 AF700 (BioLegend, Clone SK3), TIM-3 AF647 (BD Biosciences, Clone 7D3), TIGIT BV421 (BioLegend, A15153G), CTLA-4 APC/eFluor780 (eBioscience, Clone 14D3), TNF-α BV605 (BioLegend, Clone MAb11), and IFN-γ PE/Cy7 (BioLegend, Clone 4S.B3). LIVE/DEAD™ Fixable Blue Dead Cell Stain Kit (Invitrogen, ThermoFisher) was used to identify live cells. Flow cytometry was used to evaluate the production of intracellular cytokines.

### Proliferation Assays

Up to 2 x 10^7^ PBMCs or untouched CD8+ T cells were suspended in 1 ml of phosphate buffered saline (PBS, ThermoFisher) with 10% FBS. Then 5 mM of carboxyfluorescein succinimidyl ester (CFSE) solution (CellTrace™, ThermoFisher) was added, and the cells were incubated at room temperature for 8 min. The labeling reaction was quenched with 10 ml of media containing 10% FBS and cells were incubated at 37°C for an additional 10 min, then washed twice for 5 min at 300 x *g*. After CFSE staining, cells were counted with a hemocytometer via Trypan blue and cultured as indicated. CFSE replication peaks were then analyzed via flow cytometry.

### Statistical Analysis

Statistical analysis was completed by using RStudio 4.0.2 and Graph Pad Prism 8 software. Unpaired t-test, one-way ANOVA and Multiple/Pairwise Comparisons (MCPs) and Mann Whitney U test were used to compare the differences among multiple groups. A *P* value less than 0.05 was considered statistically significant.

## Results

### Inhibition of BET proteins differentially downregulates ICs on T and NK cells

To investigate the involvement of BET and AMPK proteins in IC regulation, we treated stimulated PBMCs with small molecule inhibitors. Pan-BET inhibition with JQ1 caused significant downregulation of TIM-3, PD-1 and CTLA-4 on CD4+ and CD8+ T cells, whereas TIGIT remained unchanged (Fig re 1A,B,D,E). In NK cells, JQ1 treatment significantly downregulated TIM-3 and TIGIT, whereas PD-1 and CTLA-4 remained unchanged (Figure 1C,F). Interestingly, treatment with a BRD4-selective concentration of MZ-1 (50 nM) revealed BRD4-dependent regulation of all IC markers on CD4+ T cells, with no effect on PD-1 expression in CD8+ T cells. Furthermore, BRD4 did not regulate any of these molecules on NK cells (Figure 1G-L). These findings reveal distinct patterns of IC regulation by BET proteins in T and NK cells.

**Figure 1.**
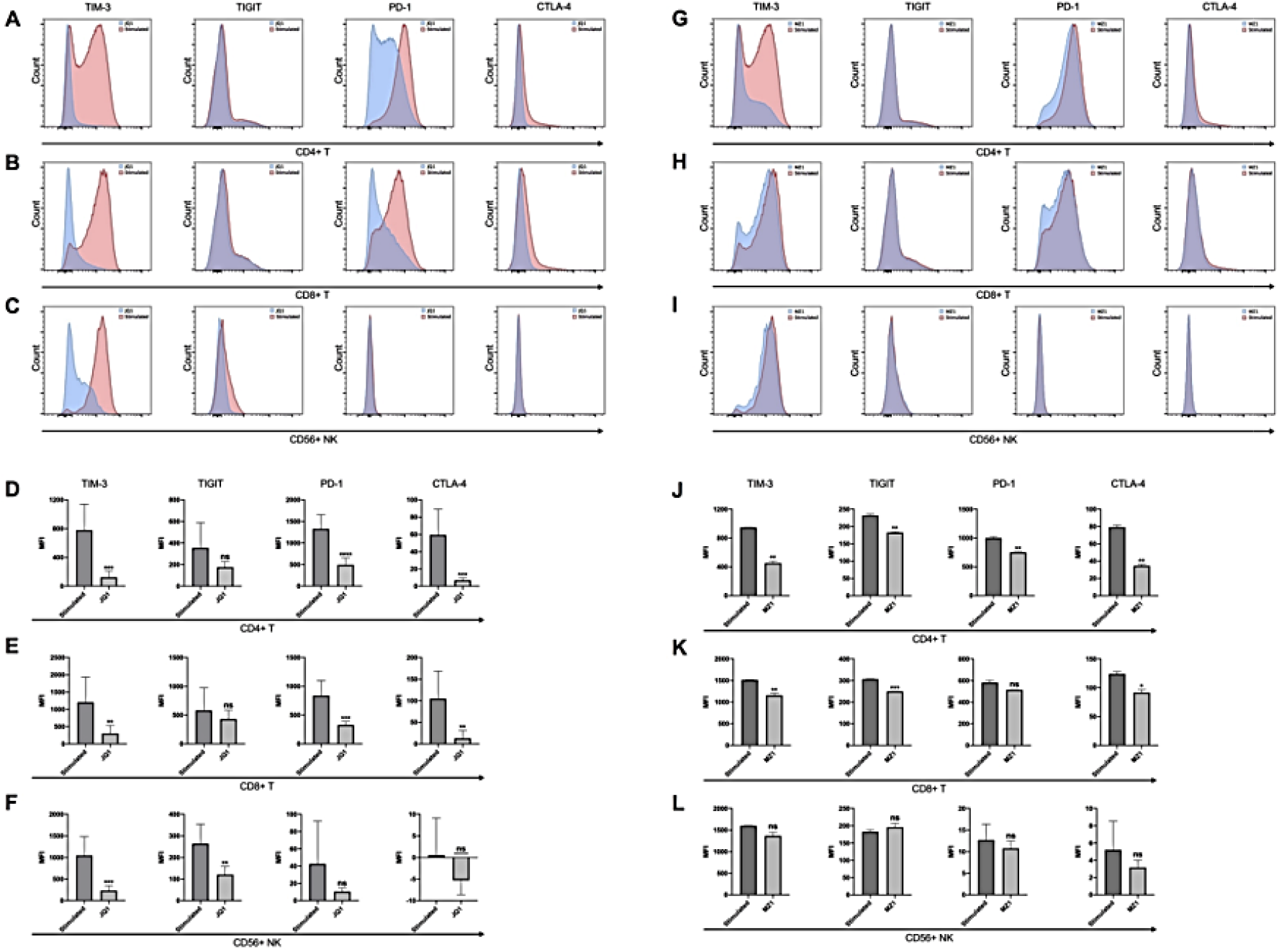
BET proteins differentially regulate TIM-3, TIGIT, PD-1 and CTLA-4 expression on both T cells and NK cells. Purified PBMCs were activated by plate-bound anti-CD3 (10 µg/ml) and soluble anti-CD28 (2 µg/ml) *ex vivo* at 37□ for 72h and treated with pan-BET inhibitor JQ1 (400 nM) or BRD4-selective PROTAC degrader MZ-1 (50 nM), respectively, for the last 48h of 72h. Cell surface fluorescence staining and flow cytometry analysis were performed to measure expression changes of TIM-3, TIGIT, PD-1, CTLA-4 on CD4+, CD8+ T cells and NK cells. (A-C, G-I) Representative histograms show expression shifts, comparing stimulated-only cells with stimulated plus JQ1 or plus MZ-1 inhibitors. X-axis: intensity of fluorescence signal. Y-axis: number of events (D-F, J-L) Corresponding bar graphs show mean fluorescence intensity (MFI) of markers on T and NK cells, comparing stimulated cells to stimulated plus JQ1 or plus MZ-1. Two-tailed t-test and Mann Whitney U test determined statistical significance. (N=6; ns, p>0.05; *, p<0.05; **, p<0.01; ***, p<0.001; ****, p<0.0001)

### Ex vivo activation of PBMCs increases expression of ICs on T and NK cells

Baseline expression of PD-1, CTLA-4, TIGIT and TIM-3 was measured via multi-parameter flow cytometry on both unstimulated control and anti-CD3/anti-CD28 stimulated CD4+ and CD8+ T cells and CD56+ NK cells (Supplementary Figure S1A). We observed significant upregulation of PD-1, CTLA-4, TIGIT and TIM-3 on activated CD4+ (Supplementary Figure S1B, S1E) and CD8+ T cells (Supplementary Figure S1C, S1F). Among CD56+ NK cells (Supplementary Figure S1D), TIM-3 showed the greatest response to stimulation compared to the other three markers (Supplementary Figure S1G), consistent with previous studies highlighting its high expression on innate cells and its further increase in NK cells upon cytokine stimulation.

### AMPK-mediated regulation of IC proteins

Treatment with the AMPK inhibitor Compound C caused significant upregulation of TIM-3 and TIGIT, but not PD-1 and CTLA-4, in CD4+ and CD8+ T cells (Figure 2A-F, H,I). Notably, NK cells also displayed similar upregulation (Figure 2G,J), indicating the existence of two distinct downstream pathways of AMPK: one regulating TIM-3 and TIGIT expression, and another regulating PD-1 and CTLA-4.

**Figure 2.**
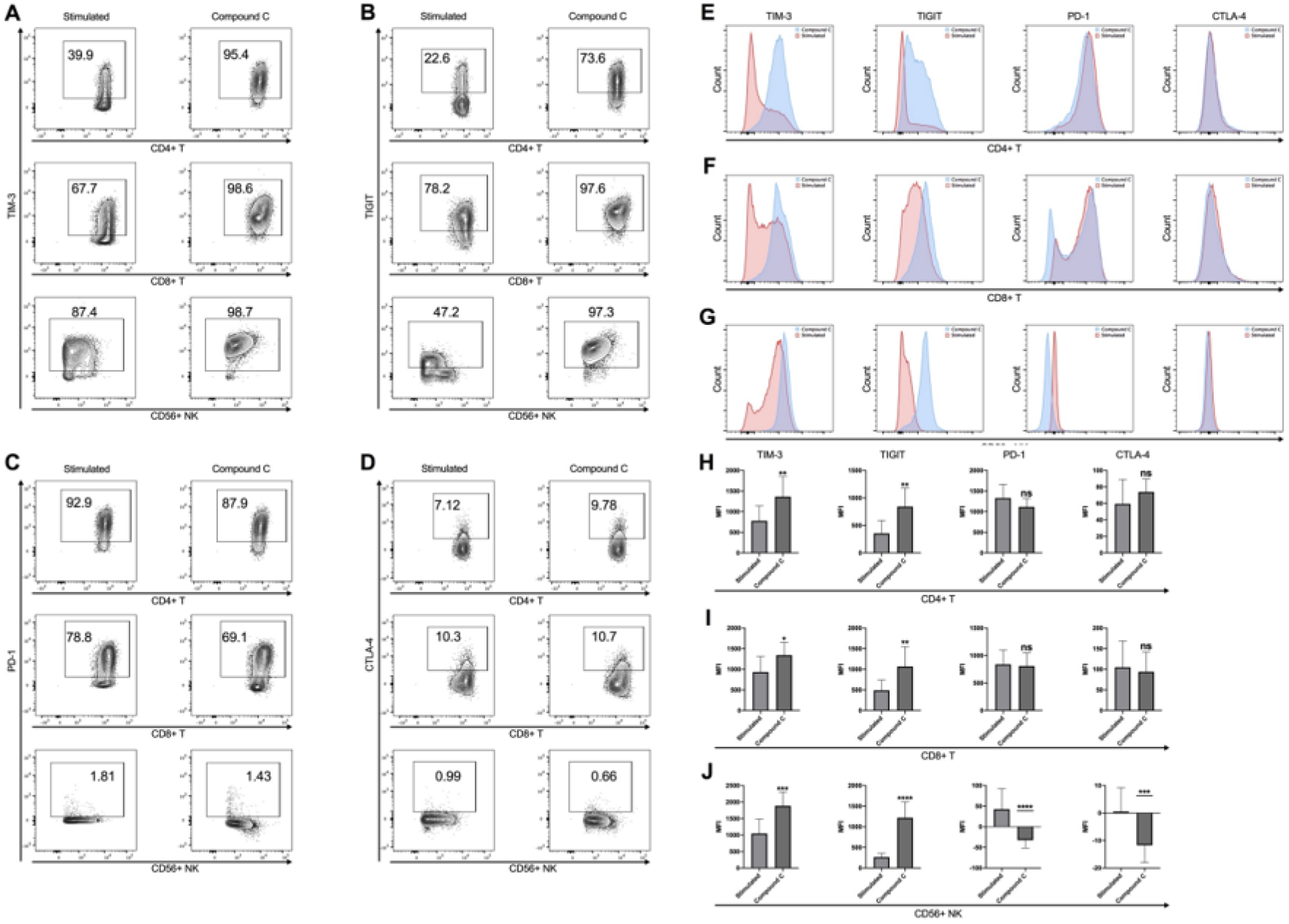
AMPK inhibition upregulates TIM-3 and TIGIT, revealing regulation that differs from PD-1 and CTLA-4, on both T cells and NK cells. PBMCs were collected and purified from normal adult donor blood. PBMCs were activated by plate-bound anti-CD3 (10 µg/ml) and soluble anti-CD28 (2 µg/ml) *ex vivo* at 37□ for 72h and treated with 2.5μM of Compound C for the last 48h of 72h. Cell surface fluorescence staining and flow cytometry analysis were performed to measure expression changes of TIM-3, TIGIT, PD-1, CTLA-4 on CD4+, CD8+ T cells and NK cells. (A-D) Contour plots show expression levels of markers in CD4+, CD8+ T cells and CD56+ NK cells, comparing stimulated to stimulated plus Compound C. (E-G) Histograms show expression shifts of each marker. X-axis: intensity of fluorescence signal. Y-axis: number of events. (H-J) Bar graphs show mean fluorescence intensity (MFI) of markers on T and NK cells, compared stimulated to stimulated plus Compound C. Two-tailed t-test and Mann Whitney U test determined statistical significance. (N=6; ns, p>0.05; *, p<0.05; **, p<0.01; ***, p<0.001; ****, p<0.0001)

### Confirmation of functional exhaustion phenotype

Given that exhaustion is a differentiation state in which cells become metabolically impaired and lose their effector functions, we validated a functional exhaustion phenotype via proliferation and cytokine assays in control experiments. Purified CD8+ T-cell populations treated with either BET or AMPK inhibitors exhibited markedly decreased replication rates and production of TNF-α and IFN-γ as indicators of effector activity, compared to control stimulated cells (Supplementary Figure S2). Inhibition of proliferation was most profound with 50 nM MZ-1, consistent with BRD4 selectivity, and the well established role of BRD4 in proliferation.

### AMPK-mediated upregulation of IC markers is not rescued by BET inhibition

To elucidate the interaction between BET proteins and the AMPK signaling pathway, stimulated cells were treated with combinations of JQ1 and Compound C or MZ-1 and Compound C. Expression of TIM-3 and TIGIT was significantly downregulated between cells treated with JQ1 alone and those with combined treatment, supporting the hypothesis of an AMPK-independent regulatory pathway. In contrast, PD-1 and CTLA-4 expression showed no significant differences between JQ1-treated and combined treatment cells, indicating an AMPK-dependent pathway (Figure 3A-F). Complex regulatory patterns were observed in cells treated with MZ-1 alone or in combination with Compound C. TIGIT exhibited an AMPK-independent pathway regulated by BRD4 in both T and NK cells. TIM-3 exhibited an AMPK-independent pathway on CD4+ T cells and an AMPK-dependent pathway on NK cells (Figure 3J,L). MZ-1 treatment resulted in the downregulation of TIM-3 on CD8+ T cells (Figure 3H). PD-1 and CTLA-4 expression clearly followed an AMPK-dependent pathway regulated by BRD4.

**Figure 3:**
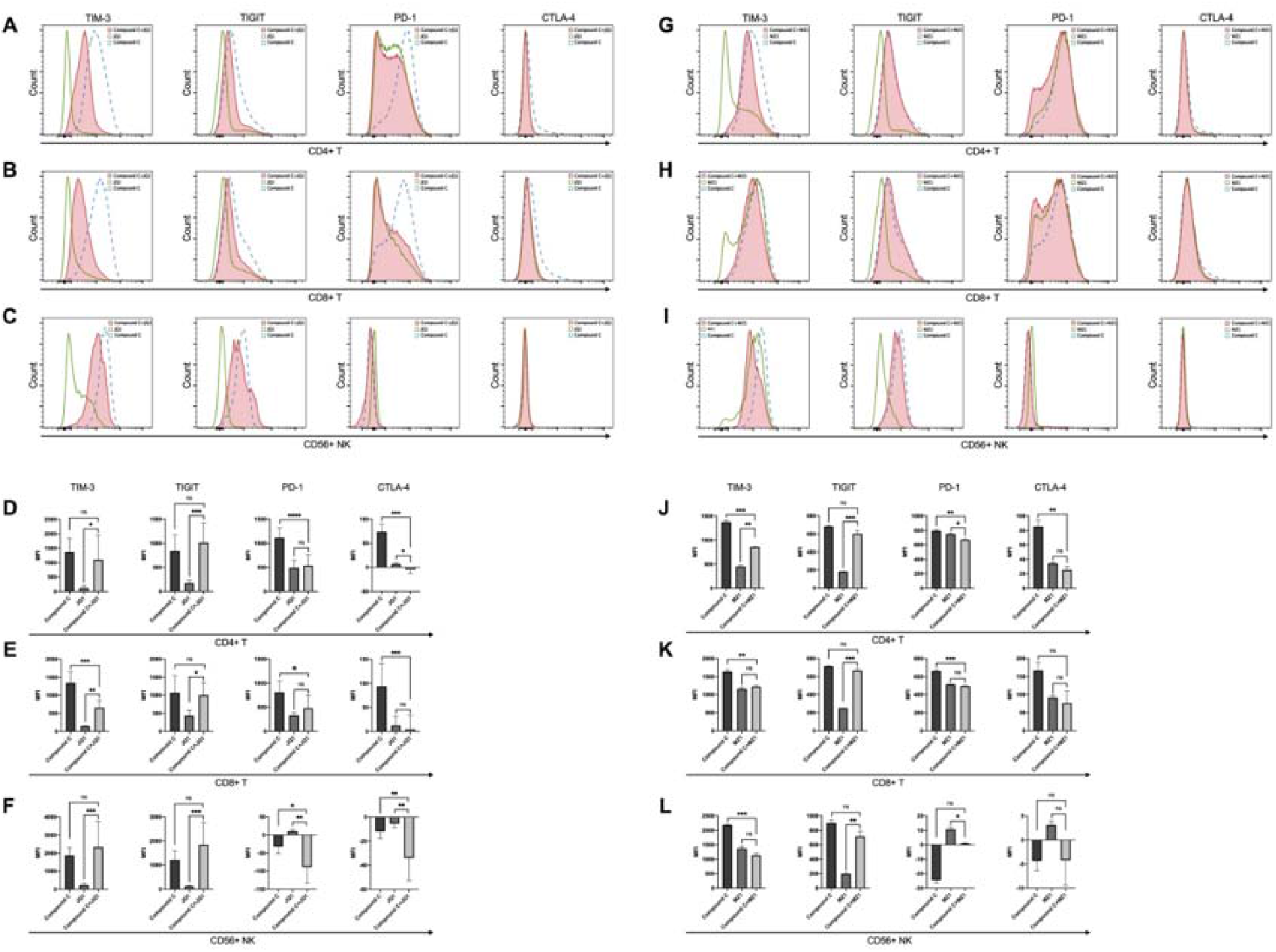
BET inhibitors do not rescue Compound C upregulation of TIM-3 and TIGIT. PBMCs were activated *ex vivo* by anti-CD3 and anti-CD28 at 37□ for 72h as above. Cell surface fluorescence staining and flow cytometry were performed to measure expression of TIM-3, TIGIT, PD-1 and CTLA-4 on both T and NK cells. Experimental groups were: stimulated plus Compound C (2.5μM), stimulated plus JQ1/MZ1 (JQ1: 400nM, MZ1: 50nM), stimulated plus Compound C and JQ1/MZ1. (A-C, G-I) Histograms combined the expression shifts of four markers in cells treated by different condition. X-axis: intensity of fluorescence signal. Y-axis: number of events. (D-F, J-L) Bar graphs show mean fluorescence intensity (MFI) of TIM-3, TIGIT, PD-1 and CTLA-4, for stimulated cells treated with Compound C, JQ1/MZ1 respectively and combined. Two-tailed t-test and Mann Whitney U test was used to measure statistical significance. (ns, p>0.05; *, p<0.05; **, p<0.01; ***, p<0.001; ****, p<0.0001)

### Proposed model of BET and AMPK regulation of IC molecules

In summary, our findings demonstrate two distinct regulatory patterns for IC markers: an AMPK-independent pathway regulating TIM-3 and TIGIT, and an AMPK-dependent pathway regulating PD-1 and CTLA-4. We propose a model illustrating the interaction between BET proteins and AMPK signaling in the regulation of these markers (Supplementary Figure S3). The upregulation of TIM-3 and TIGIT by Compound C suggests the existence of an unknown downstream pathway that is normally inhibited by activated AMPK. BET proteins may act as coregulators of AMPK-regulated PD-1 and CTLA-4 gene expression, while displaying a separate regulation pathway for TIM-3 and TIGIT.

## Discussion

We explore regulatory networks of BET proteins and AMPK signaling that regulate immune checkpoint expression in human primary T and NK cells. We find that BET proteins regulate PD-1 and CTLA-4 through an AMPK-dependent pathway and TIM-3 and TIGIT through an AMPK-independent pathway. Better understanding of this network is important because many solid tumors occur in patients who have comorbid metabolic disease, particularly Type 2 diabetes, and have worse outcomes. The major first-line therapy for T2D is metformin, which is well established as an activator of AMPK. However, the regulatory networks relevant for immune checkpoint blockade in these patients have not been studied. This knowledge gap reveals an urgent public health challenge, as more than 120 million Americans are currently diagnosed with diabetes or pre-diabetes, 90% of which is T2D. Yet should they develop cancer, under the current standard of care in medical oncology, they are not treated differently in any significant way from metabolically normal cancer patients, including for recommendations for ICB therapies.

Much research effort has been invested to understand the mechanisms of action of BET bromodomain family of transcriptional coregulators, comprised of BRD2, BRD3 and BRD4 in somatic cells,^9^ which play a critical role in tumor cell proliferation for a wide array of solid tumor types, as well as leukemias and lymphomas.^10-15^ This family is also recently appreciated to be important for tumor immune infiltrate function, especially CD8+ T cell effectors, and expression^17,18,19^ of PD-1 and PD-L1, which are critical immune checkpoint genes that have been leveraged as novel targets for ICB in several cancers,^18^ including TNBC.^7,8,13,15,17^

NK cells also play a pivotal role in cancer immunity due to their unique capability to detect and eliminate malignant cells independent of prior sensitization.^3^ This intrinsic ability to exert cytotoxicity, secrete crucial cytokines, and even modulate the adaptive immune response underscores their importance in thwarting tumor progression, curtailing metastatic spread, and counteracting the evasive tactics employed by cancer cells. Currently, no study has indicated the regulation of immune checkpoints in NK cells by BET proteins. Our results demonstrate that NK cells share a similar regulation pattern with checkpoint genes in T cells, which can be manipulated via small molecule inhibitors.

The significance of the AMPK pathway in cellular metabolism takes a new dimension as here we unveil its intricate involvement in the regulation of immune checkpoints. Our study uniquely highlights not only the influence of this pivotal pathway on checkpoint ligands but also the distinct regulatory patterns governing different molecules. These findings open promising avenues for the development of innovative therapies that target specific expression profiles. This advance holds particular importance in cases of AMPK dysfunction such as in T2D, the most common metabolic disorder that is also known to increase cancer risk. This disease is commonly managed with metformin, a drug that modulates AMPK; however, there is conflicting evidence whether metformin is protective or harmful for tumor progression, underscoring the need for comprehensive characterization to unravel the AMPK pathway’s intricate interplay in individualized patient biology.

The tumor microenvironment of breast cancer patients with comorbid T2D is entirely different from similar breast cancer patients who lack metabolic complications. In particular, the well-known immune abnormalities of T2D patients alter the tumor microenvironment and likely contribute to failure of anti-tumor immunity and enable distant metastases. We recently modeled the microenvironment of ER+ breast cancer in T2D and observed the emergence of immunosuppressive tumor infiltrating lymphocytes.^30^ It is unknown how this subset responds to AMPK activators such as metformin. We suggest that more detailed and personalized profiling of metabolic features and microenvironment characteristics in TNBC patients with comorbid T2D will enable development of better clinical approaches to reduce risk for metastasis and mortality, including improved responses to ICB therapy.

Likewise, individual BET proteins demonstrate complex and sometimes opposing regulation of key genes that are critical for tumor progression, including epithelial-to-mesenchymal transition.^15^ Similarly, individual IC genes such as PD-1 are regulated by BRD2 and BRD4, but not BRD3,^17^ which implies that personalized profiling of the specific tumor type and tumor infiltrating immune cell is required to justify a particular ICB strategy combined with a selective BET inhibitor.

A limitation of this work is that the PBMCs we profiled were purified from normal adult donors. PBMCs from a range of T2D patients, grouped by age, race, sex, and duration of T2D, as well as the effectiveness of glucose control with medications, will be important to understand how the regulatory network is altered in a personalized fashion under the chronic stimulation, immune exhaustion and unresolved inflammation that is characteristic of most T2D patients. These networks are also likely to be affected by the stage and type of cancer for which the patient is being considered for ICB therapy, as well as cancer-specific treatments, such as radiation, cytotoxic chemotherapy or targeted agents. The responsiveness of the AMPK network in the tumor immune infiltrates may also differ from circulating PBMCs, which we used in this initial study. Nevertheless, our findings here strongly advocate for personalized profiling of patient ICB regulatory networks before a specific agent, ICB monoclonal antibody, whether against PD-1, PD-L1, CTLA-4, or other targets including TIM-3 and TIGIT, and combinations with a BET inhibitor or related drug, can be well justified for any cancer patient.

In summary, we reveal unexpected interactions among BET protein family, the AMPK signaling pathway, and immune checkpoint expression. Our findings contribute to the growing understanding of immune checkpoint regulation and hold potential for the development of targeted therapies to combat immune exhausted and related diseases, particularly breast cancer in patients who have comorbid T2D.

## Acknowledgments

We thank Boston University-Boston Medical Center Cancer Center faculty R. Flynn, N. Ganem and V. Perissi, for helpful comments and suggestions. We thank Anna Belkina, Shari Brezinsky and Brian Tilton, the BUMC Flow Cytometry Core Facility, Matthew Bo Au of the BUMC Analytical Core Facility. This work was supported by the Cancer Moonshot and Cancer Systems Biology Consortium of the National Cancer Institute (G.V.D.: U01CA182898, U01CA243004 and R01CA222170)

## Author Contributions

CRediT: Christina S. Ennis: Conceptualization, Investigation, Methodology, Writing – original draft; Kunlin Huang: Conceptualization, Investigation, Methodology, Writing – original draft; Allison N. Casey: Investigation, Methodology; Michael Seen: Investigation, Clinical coordination, Methodology; Gerald V. Denis: Funding acquisition, Project administration, Resources, Conceptualization, Project administration, Writing – original draft, Writing – review & editing.

## Data availability statement

The data supporting this study are available from the corresponding authors upon reasonable request.

## Supplementary Data

Supplementary information is available in a separate file, comprising Supplementary Figures S1-S3.

