## Supplementary Figures S1-S3 for "Unraveling AMPK and BET regulation of immune checkpoint biology: implications for personalized medicine"

**Supplementary Information**

**Supplementary Data and Figures**

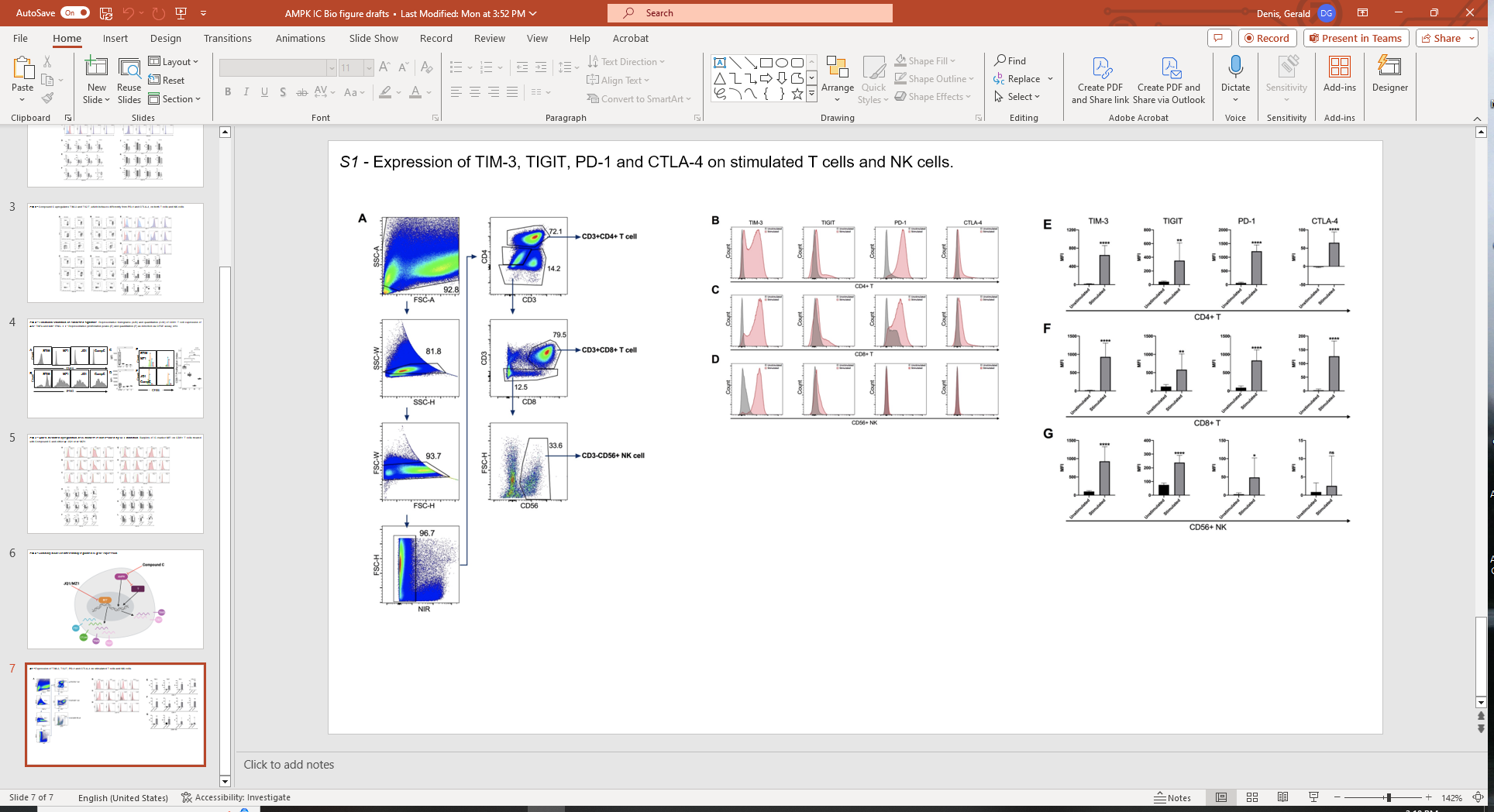
**Supplementary Figure S1**

**Supplementary Figure S1.** PBMCs were collected and purified from normal adult donor blood, and were activated by plate-bound anti-CD3 (10 µg/ml) and soluble anti-CD28 (2 µg/ml) *ex vivo* at 37℃ for 72h. Fluorescence cell surface staining and flow cytometry were performed to analyze the expression of immune checkpoint proteins. (A) Gating strategies used to identify cell types, i.e. CD4+ T cells, CD8+ T cells and CD56+ NK cells. (B-D) Representative histograms show expression changes of these immune checkpoints on both T and NK cells. Expression shifts of immune checkpoints on T and NK cells are shown by histogram, comparing unstimulated cells to stimulated. The X-axis refers to the intensity of the fluorescence signal, and the Y-axis refers to the number of events. (E-G) Bar graphs show mean fluorescence intensity (MFI) of the markers on T and NK cells, comparing unstimulated cells to stimulated cells. Two-tailed unpaired t-test and Mann Whitney U test determined statistical significance. (N=6; ns, p>0.05; *, p<0.05; **, p<0.01; ***, p<0.001; ****, p<0.0001)

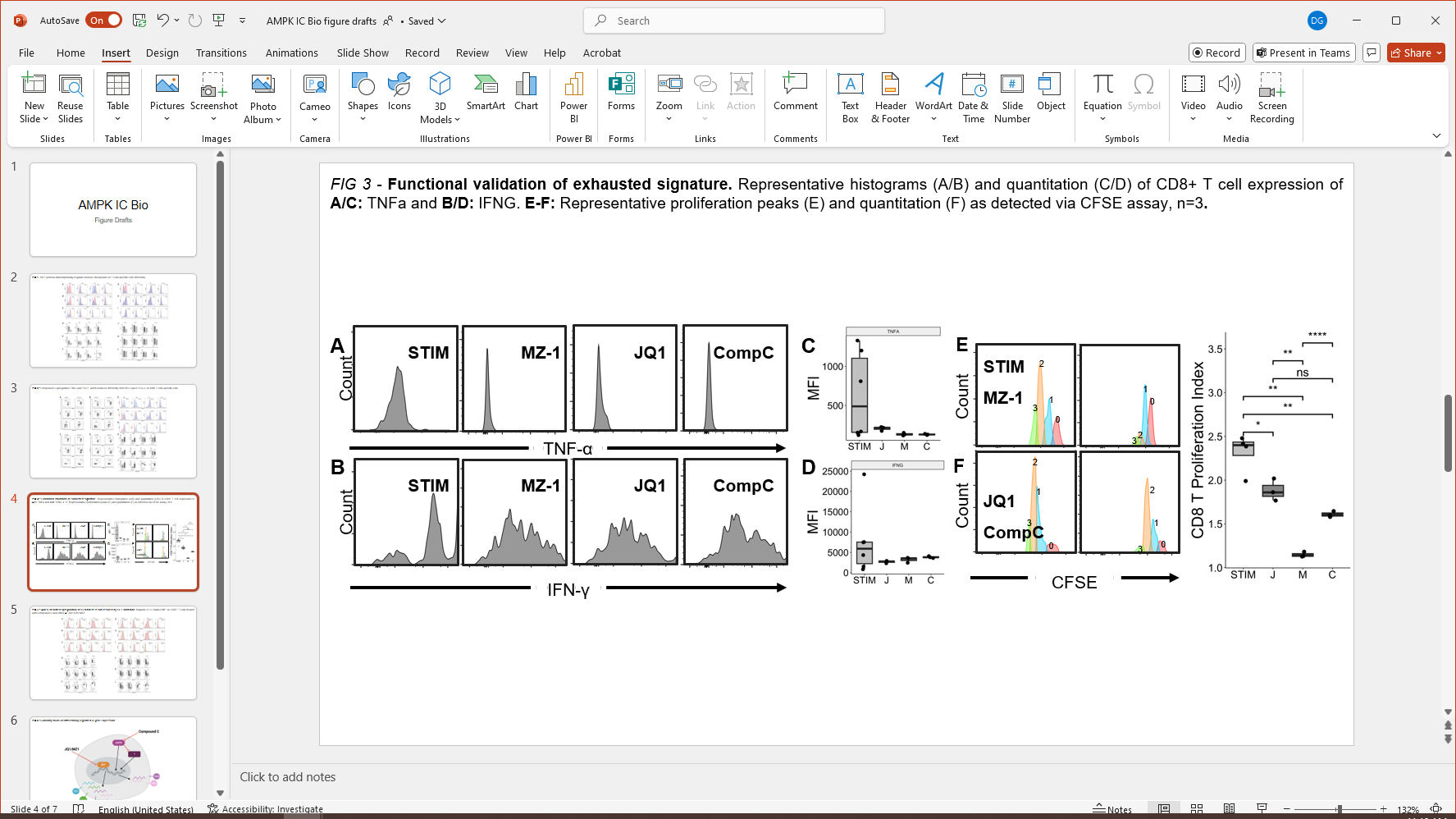
**Supplementary Figure S2**

**Supplementary Figure S2.** Functional validation of exhausted signature. Representative histograms (A,B) and quantitation (C,D) of CD8+ T cell expression of A,C: TNF-α; and B,D: IFN-γ. E,F: Representative proliferation peaks (E) and quantitation (F) as detected via CFSE assay, n=3. PBMCs were purified from normal adult donor blood, activated and analyzed as described in Supplementary Figure S1. Two-tailed unpaired t-test and Mann Whitney U test determined statistical significance. (ns, p>0.05; *, p<0.05; **, p<0.01; ***, p<0.001; ****, p<0.0001) STIM, activation by plate-bound anti-CD3 (10 µg/ml) and soluble anti-CD28 (2 µg/ml); J, STIM plus JQ1 (400 nM); M, STIM plus BRD4-selective PROTAC degrader MZ-1 (50 nM); C, STIM plus Compound C (2.5μM).

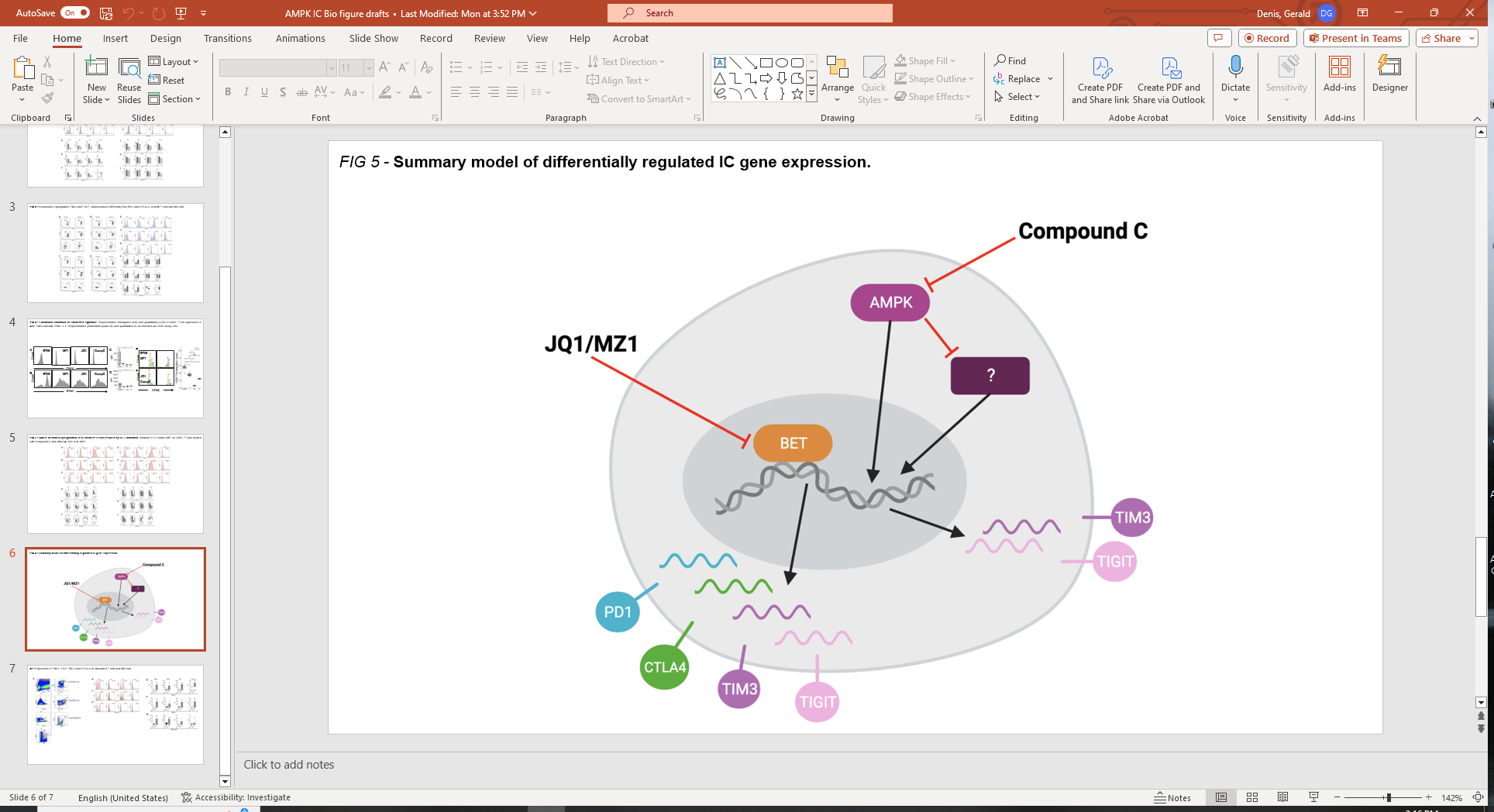

**Supplementary Figure S3.** Model for BET and AMPK regulation of TIM-3, TIGIT, PD-1 and CTLA-4. Regulation of immune checkpoints by BET proteins and AMPK are both independent and dependent. BET proteins act as co-transcriptional regulators for multiple genes but also function as one of the downstream regulators of AMPK and contribute to expression of PD-1 and CTLA-4. When compound C initiates an unknown (?) pathway and upregulates expression of TIM-3 and TIGIT, BET effectors do not participate in that regulation, which means that BET proteins are not a cofactor of the new pathway to regulate TIM-3 and TIGIT expression. Conversely, the BET proteins themselves use a regulatory pathway independent of AMPK, which regulates TIM-3 and TIGIT.
